# Clustering Analysis to Explore Cohorts in Comorbid Patients

**DOI:** 10.1101/396481

**Authors:** Rasika Karkare

## Abstract

Multimorbidities are associated with significant burden on the healthcare system and the lack of accurate and pertinent statistical exploratory techniques have often limited their analysis. Here we employ exploratory hierarchal agglomerative clustering (HAC) of multimorbidities in the inpatient population in the state of Ohio. The examination exposed the presence of ten discrete, clinically pertinent groups of multimorbidities within the Ohio inpatient population. This method offers an assessable empirical exploration of the multimorbidities present in a specific geographic populace.

## 2. INTRODUCTION

Patients with more than one chronic conditions are referred to as multimorbid patients. The incidence of such multimorbid patients has increased due to the rise in the aging population in the US leading to significant impact on the overall health care outcomes. Numerous mechanisms may lie beneath these multimorbidities, including direct relationship, associated risk features, heterogeneity, and individual differences ^1,2^. There has been an augmented acknowledgment of its bearing and the significance of improving consequences for such multimorbid patients^3–5^. The lack of proper and statistical exploratory techniques have often limited the analysis of such chronic multimorbidities.

Clustering, is an unsupervised data mining algorithm that groups similar units into similar clusters, thus partitioning dissimilar objects into other clusters^6–10^. Hierarchical agglomerative clustering is a repeated subdivision of a dataset into clusters at an progressively finer granularity^11^. HAC has previously been employed to find clusters in microarray data^12^, employed in nursing research^10^, and to find molecular genetic markers^9^. In the current study, we use unsupervised clustering exploration to discover the cohorts of cohorts present in the Ohio patient population.

## 3. METHOD

### 3.1 DATA

Publicly available data from the Ohio Department of Health (https://www.odh.ohio.gov/) was used in this study. The data is devoid of all PPI and PHI and follows state and national HIPPA standards. Several exclusions including pregnant females, cancer, patients in hospice or long term care were excluded from the analysis. Further, patients only within the ages of 40 to 80 were identified for this multimorbidity analysis. Identification of conditions within unit associates were founded on an in/outpatient data matched to *International Classification of Diseases (ICD-11)* codes^13^

### 3.2 DATA EXPLORATION

Apache Hadoop database was used to store, query and extract the dataset extracted from the online resource. One step hot encoding was used for dichotomous target features for selected disorders. Demographic variables were appended to the data set to enable broader interpretation. Data missing at random was imputed by median/mode for continuous/categorical features.

### 3.3 STATISTICAL ANALYSIS

The R suite of statistical programs was used for quantitative analysis (Version 0.9). Statistical procedures used are described previously in other studies ^6,14–20^.

### 3.4 CLUSTERING ALGORITHM

A bottom up clustering approach also known as Agglomerative Hierarchical Clustering (AHC) was used for this analysis. We assumed monotonicity of the operation and the number of clusters was determined by subject matter expertise and medical interpretability. The resulting analysis was visualized using tree-maps or dendrograms. The algorithm employed Gower’s matrix for similarity distance computations ^21,22^.

## 4. RESULTS

### 4.1 HAC DIVULGES TEN MULTIMORBID CLINICALLY RELEVANT CLUSTERS

HAC showed the presence of ten multimorbid cohorts in the patient population (Figure 1). Overall, we saw a higher proportion of females in the data (58.3 % females). The median age in the dataset was 73.0 years.

**Figure 1:**
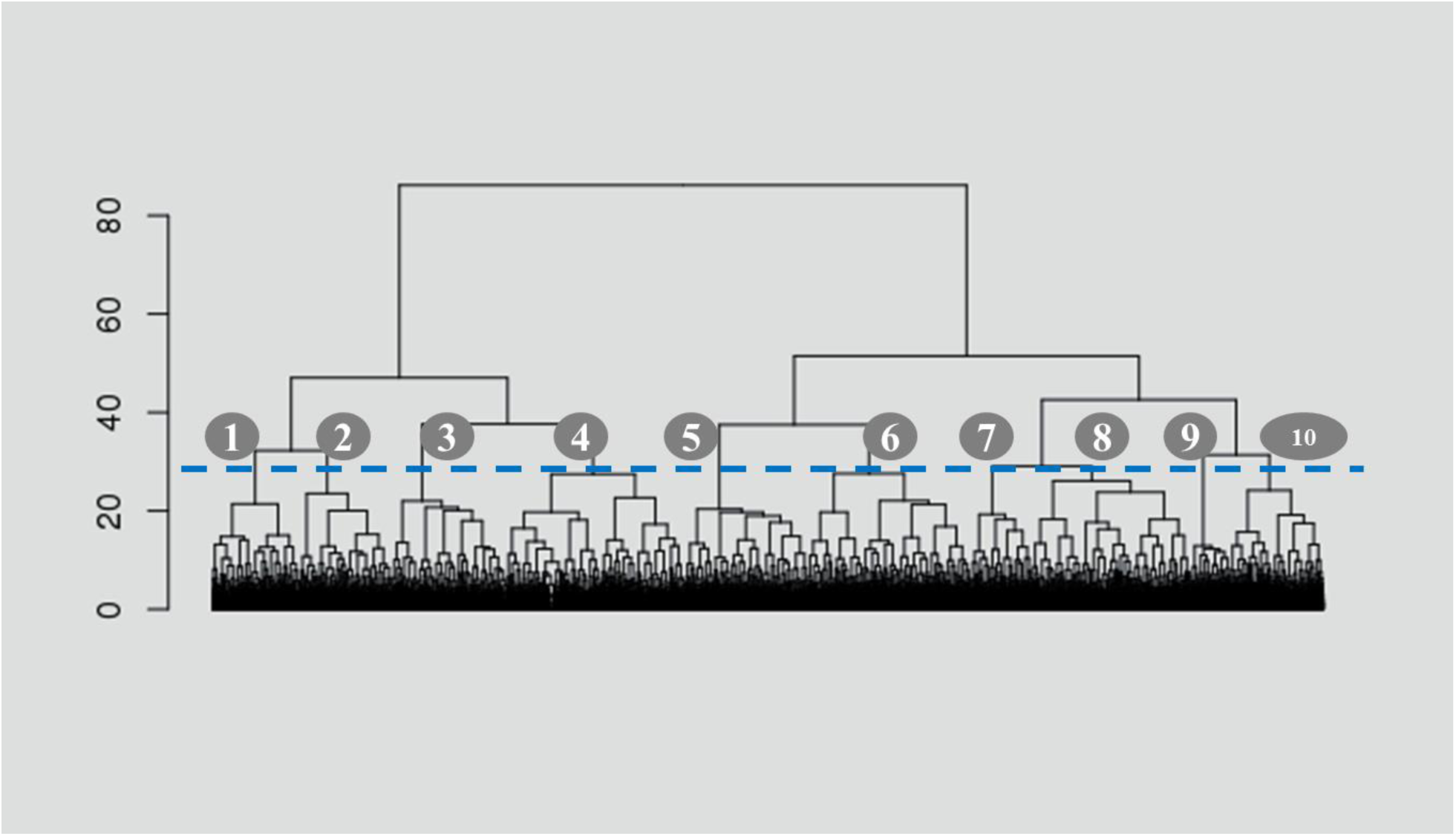
HAC presents nine separate cohorts in the patient data (tree cut at height=27; broken line).

Quantitative and qualitative descriptions of the clusters showed overall homogeneity (Table 1). anWe summarized out cluster summaries based on subject matter expertise (Table 2) as follows: metabolic syndromes (cluster 1), cancer (cluster 2), gastro related diseases (cluster 3), hypertensive patients (cluster 4), stomach cancer and related conditions (cluster 5), heart failure and COPD (cluster 6), Obesity related morbidities like lower back pain (cluster 7), Epilepsy (cluster 8), osteoarthritis (cluster 9) and old age related disorders like Parkinson’s (cluster 10). Other patient specific metrics were available but not included in the study due to PII restrictions.

**Table 1:** Condition prevalence by cluster/cohort. The age (years, median), gender makeup (%) and the patient prevalence is shown for each HAC cluster.

| Cluster | 1 | 2 | 3 | 4 | 5 | 6 | 7 | 8 | 9 | 10 |
| --- | --- | --- | --- | --- | --- | --- | --- | --- | --- | --- |
| size | 1193 | 2319 | 1600 | 2108 | 1139 | 1494 | 1212 | 2065 | 807 | 507 |
| age (mean) | 68.5 | 62.2 | 75.7 | 76.4 | 77 | 72.9 | 69.9 | 77.5 | 79.2 | 81.1 |
| males (%) | 42.5 | 45.2 | 57.7 | 48.8 | 40.9 | 56.5 | 32.6 | 59.3 | 48.9 | 43 |
| females (%) | 57.5 | 54.8 | 42.3 | 51.2 | 59.1 | 43.5 | 67.4 | 40.7 | 51.1 | 57 |
| Hospital Visits (mean) | 0.73 | 0.94 | 1.14 | 0.98 | 0.68 | 0.89 | 0.94 | 0.91 | 0.82 | 1.07 |
| Diabetes Mellitus (%) | 0 | 0.2 | 0.2 | 0.1 | 0.2 | 0.2 | 0.2 | 0 | 0 | 0.2 |
| Heart Failure (%) | 0.4 | 0.4 | 0.4 | 0.4 | 0.2 | 0.7 | 0.2 | 0.2 | 0.5 | 0.2 |
| COPD (%) | 0.2 | 0.6 | 0.3 | 0.4 | 0.5 | 1.5 | 0.1 | 0.2 | 0.4 | 0.2 |
| Ischemic Heart Disease (%) | 0 | 0.2 | 0 | 0 | 0.1 | 0.4 | 0 | 0.1 | 0 | 0 |
| Otitis Media (%) | 63.4 | 16.4 | 83 | 75.7 | 30.5 | 67.9 | 49.6 | 28.2 | 29.5 | 44.4 |
| Peptic Ulcer Disease (%) | 19.4 | 4.7 | 81.9 | 71.5 | 13.3 | 34.1 | 44.6 | 53.9 | 49.4 | 50.3 |
| Hypertension (%) | 32.7 | 23.8 | 35.4 | 67.3 | 25.1 | 23 | 66.5 | 46.9 | 47.8 | 45.6 |
| Epilepsy (%) | 22 | 6.7 | 77.6 | 67.1 | 28.4 | 14.3 | 57.9 | 78.3 | 33.2 | 62.7 |
| Chronic Thyroid Disorders (%) | 1.3 | 1 | 1 | 1.5 | 1.1 | 0.5 | 2.9 | 1.2 | 0.7 | 1.2 |
| Depression (%) | 3.1 | 2.8 | 2.4 | 6.3 | 2.5 | 1.9 | 4.7 | 2.9 | 3.5 | 4.5 |
| Cholelithiasis Cholecystitis (%) | 95.3 | 51.5 | 99 | 98.3 | 93.4 | 88 | 93 | 95.7 | 93.9 | 97 |
| Iron Deficiency Anemia (%) | 1.8 | 7.7 | 4.6 | 3.8 | 1.2 | 3.3 | 7.3 | 5 | 5.1 | 11.8 |
| Osteoarthritis (%) | 31.3 | 17.4 | 24.9 | 34.3 | 28.4 | 24.2 | 33.9 | 26.1 | 37.2 | 25.6 |
| Rheumatoid Arthritis (%) | 23.7 | 32.9 | 20.9 | 38.5 | 24.5 | 13.4 | 70.1 | 17.5 | 18.3 | 71 |
| Colorectal Cancer (%) | 2.6 | 2.5 | 2.9 | 4.4 | 1.8 | 2.3 | 3.1 | 2.3 | 2.9 | 3.4 |
| Lung Cancer (%) | 10.7 | 6.3 | 23.1 | 15.8 | 12.2 | 10.8 | 11.5 | 9.2 | 16 | 15.2 |
| Hyperlipidemia (%) | 60.9 | 17.9 | 27.9 | 49.8 | 89.5 | 14 | 61.6 | 24.3 | 42.5 | 42.2 |
| Cerebrovascular Disease (%) | 4.9 | 2.5 | 2.6 | 5.6 | 9 | 2.4 | 13.7 | 2.4 | 4.8 | 3.9 |
| Headaches Migrane Othrs (%) | 1.4 | 4.4 | 3.3 | 3.6 | 3.7 | 3.1 | 1.3 | 2.5 | 5 | 2.4 |
| Chronic Renal Failure (%) | 0.8 | 3.8 | 1.8 | 2.6 | 1.6 | 2.2 | 3.2 | 5.2 | 3.8 | 2.2 |
| Kidney Stones (%) | 77.9 | 44.8 | 91.2 | 92.4 | 72.5 | 69.5 | 84.4 | 84.3 | 79.3 | 88.8 |
| Diverticular Disease (%) | 14.6 | 11.1 | 33.4 | 35.1 | 11.4 | 14.9 | 26.3 | 46.4 | 36.6 | 64.5 |
| Low Back Pain (%) | 7.7 | 15 | 4.8 | 8.2 | 4.5 | 4.2 | 28 | 7.1 | 6.8 | 18.1 |
| Nonspec Gastr Dyspepsia (%) | 32.1 | 7.5 | 79.3 | 78.9 | 23.4 | 35.4 | 18.2 | 29.9 | 36.4 | 49.5 |
| Sickle Cell Anemia (%) | 3.3 | 4 | 4.8 | 7 | 4.7 | 3.7 | 6.3 | 3.5 | 5.2 | 3.2 |
| Multiple Sclerosis (%) | 6.1 | 4.6 | 4.4 | 13.4 | 11.4 | 2.2 | 11.8 | 5.5 | 12.5 | 11.2 |
| Inflammatory Bowel Disease (%) | 56.4 | 39.5 | 22.5 | 60.8 | 48.3 | 14.3 | 72.5 | 39.4 | 40.3 | 44.2 |
| Hemo Congntl Coagulopathies (%) | 63.4 | 44.2 | 50.4 | 68.6 | 54.4 | 31.9 | 75.5 | 50.8 | 52.8 | 76.1 |
| Systemic Lupus Ery (%) | 0.2 | 0.6 | 0.3 | 0.8 | 0 | 0.1 | 0.2 | 0 | 0.4 | 0.2 |
| Prostate Cancer (%) | 0.2 | 1.1 | 0.2 | 0.5 | 0.4 | 0.4 | 0.9 | 0.3 | 0.4 | 0.2 |
| Ovarian Cancer (%) | 2.3 | 5.1 | 0.5 | 1.2 | 1.5 | 1.4 | 2.8 | 1.4 | 2 | 1.4 |
| Endometrial Cancer (%) | 1.4 | 1.1 | 0.2 | 1.2 | 0.4 | 0.6 | 3.3 | 0.6 | 0.4 | 0.8 |
| Cervical Cancer (%) | 5.1 | 2.1 | 7.7 | 7.5 | 6 | 5.8 | 2 | 7 | 8.6 | 9.5 |
| Hodgkin Dis Lymphoma (%) | 0.1 | 1 | 0.1 | 0.3 | 0.4 | 0.1 | 0 | 0.2 | 0.5 | 0.2 |
| Leukemia Myeloma (%) | 0.3 | 0.9 | 0.2 | 0.5 | 0.3 | 0.5 | 0.7 | 0.2 | 0.5 | 0.2 |
| Malignant Melanoma (%) | 0.2 | 0.5 | 0 | 0.2 | 0.3 | 0.1 | 0.1 | 0 | 0.1 | 0.2 |
| Head Neck Cancer (%) | 1.1 | 3 | 1.4 | 1.2 | 1.5 | 1.7 | 1.9 | 1.7 | 1.4 | 1.4 |
| Esophageal Cancer (%) | 1.3 | 2.5 | 2.4 | 2.3 | 1.8 | 3.4 | 0.8 | 1.8 | 3.5 | 1.6 |
| Stomach Cancer (%) | 1.3 | 1.1 | 1.3 | 1 | 3.8 | 1 | 1.3 | 3.1 | 2 | 1 |
| Pancreatic Cancer (%) | 0.3 | 2.1 | 0.4 | 0.6 | 1.1 | 0.4 | 0.3 | 1.2 | 0.4 | 0.2 |
| Pancreatitis (%) | 0.2 | 0.6 | 0.1 | 0.1 | 0 | 0.5 | 0.2 | 0.3 | 0.6 | 0.2 |
| Hepatitis (%) | 2.5 | 6.9 | 2 | 4.7 | 1.8 | 1.5 | 3.6 | 1.4 | 2.5 | 2.8 |
| Peripheral Artery Disease (%) | 2.8 | 5.5 | 1.8 | 1.9 | 2.2 | 4.5 | 4.3 | 1.7 | 1.6 | 2.2 |
| Endometriosis (%) | 15.5 | 8.3 | 40.9 | 86 | 24.1 | 12.4 | 44.8 | 38.9 | 41.6 | 60.2 |
| Ventricular Arrhythmia (%) | 0.1 | 0.5 | 0 | 0 | 0 | 0 | 0.2 | 0 | 0 | 0 |
| Lyme Disease (%) | 2.2 | 1.5 | 12.2 | 8.3 | 2.2 | 3.1 | 5.9 | 10.4 | 5.3 | 3.7 |
| Female Infertility (%) | 0 | 0.1 | 0.1 | 0 | 0.2 | 0.1 | 0.2 | 0 | 0.1 | 0 |
| Menopause (%) | 0.1 | 0.1 | 0 | 0 | 0 | 0 | 0 | 0 | 0 | 0 |
| Glaucoma (%) | 3.7 | 2.8 | 1.5 | 3.7 | 4.7 | 1.7 | 10.1 | 2.7 | 3.5 | 4.1 |
| Low Vision Blindness (%) | 9.3 | 3.8 | 17.6 | 22.2 | 9.2 | 10.9 | 14.4 | 10.7 | 52.3 | 31.2 |
| Cataract (%) | 0.6 | 0.6 | 1.7 | 1.5 | 1.3 | 0.9 | 0.9 | 0.5 | 0.4 | 3.2 |
| Other Cancer (%) | 11.7 | 5.2 | 13.9 | 43.1 | 13.9 | 11.1 | 24.1 | 13.3 | 64.6 | 30.4 |
| Dementia (%) | 3.2 | 11.2 | 4.4 | 6.4 | 7.1 | 10.1 | 4.5 | 7.2 | 7.4 | 4.3 |
| Osteoporosis (%) | 5.5 | 4.7 | 9.5 | 11.5 | 8.6 | 16.9 | 10.6 | 8.2 | 10.7 | 86.8 |
| Obesity (%) | 7.3 | 6.6 | 7 | 16 | 20.8 | 2.9 | 24.8 | 7.1 | 9 | 21.3 |
| Oral Cancer (%) | 87.1 | 11 | 43.7 | 59.3 | 11.8 | 17.3 | 62.4 | 18.1 | 20.3 | 21.3 |
| Cystic Fibrosis (%) | 0 | 1.7 | 0.4 | 0.5 | 0.6 | 0.2 | 0.3 | 0.6 | 0.2 | 0.2 |
| Neurosis (%) | 0.2 | 0.3 | 0 | 0 | 0 | 0 | 0.2 | 0.1 | 0 | 0 |
| Psychoses (%) | 1.3 | 5.5 | 1.9 | 1.8 | 1.1 | 1.3 | 6.7 | 1.7 | 1.5 | 8.3 |
| Eating Disorders (%) | 0 | 0.2 | 0.1 | 0.1 | 0 | 0 | 0.2 | 0 | 0 | 0.6 |
| Disrupt Childhd Disorders (%) | 0.2 | 0.1 | 0.1 | 0 | 0 | 0 | 0.2 | 0 | 0 | 0.2 |
| Substance Reltd Disorders (%) | 7 | 19.6 | 4.4 | 22 | 5.4 | 2.9 | 28.1 | 5.3 | 3.8 | 7.9 |
| Skin Cancer (%) | 4.9 | 3.8 | 5.8 | 5.8 | 19.5 | 5.6 | 7 | 21.4 | 11 | 9.1 |
| Congenital Heart Disease (%) | 0.8 | 0.4 | 1.5 | 1.6 | 0.4 | 0.5 | 0.9 | 1.1 | 1.5 | 1.4 |
| Periodontal Disease (%) | 0.8 | 0.7 | 0.6 | 0.4 | 1.1 | 0.7 | 0.4 | 0.4 | 0.4 | 0.6 |
| Chronic Fatigue Syndrome (%) | 0.6 | 0.6 | 0.6 | 0.9 | 0.4 | 0.2 | 1.6 | 1.1 | 0.6 | 0.6 |
| Fibromyalgia (%) | 3.9 | 3.8 | 1.2 | 2.7 | 3.1 | 0.7 | 14.9 | 1.2 | 2.2 | 2 |
| Parkinson Disease (%) | 1.3 | 0.9 | 1.6 | 2.8 | 2.3 | 2.7 | 2.1 | 1.9 | 1.7 | 15.4 |
| Hypercoaguable Syndome (%) | 1 | 1.1 | 0.8 | 1.2 | 1.3 | 1.2 | 1.7 | 0.9 | 1.4 | 1.4 |
| Post Partum BH Disorder (%) | 0 | 0.2 | 0 | 0 | 0 | 0 | 0 | 0 | 0 | 0 |
| Metabolic Syndrome (%) | 23.3 | 6.8 | 3.3 | 3.2 | 3.6 | 2.5 | 7 | 5.7 | 4.8 | 4.3 |
| Psyc Dis rltd Med<br>Condtns (%) | 1.5 | 2.9 | 1.5 | 2.2 | 1.6 | 2 | 4.3 | 1.3 | 2.9 | 23.1 |

**Table 2:**
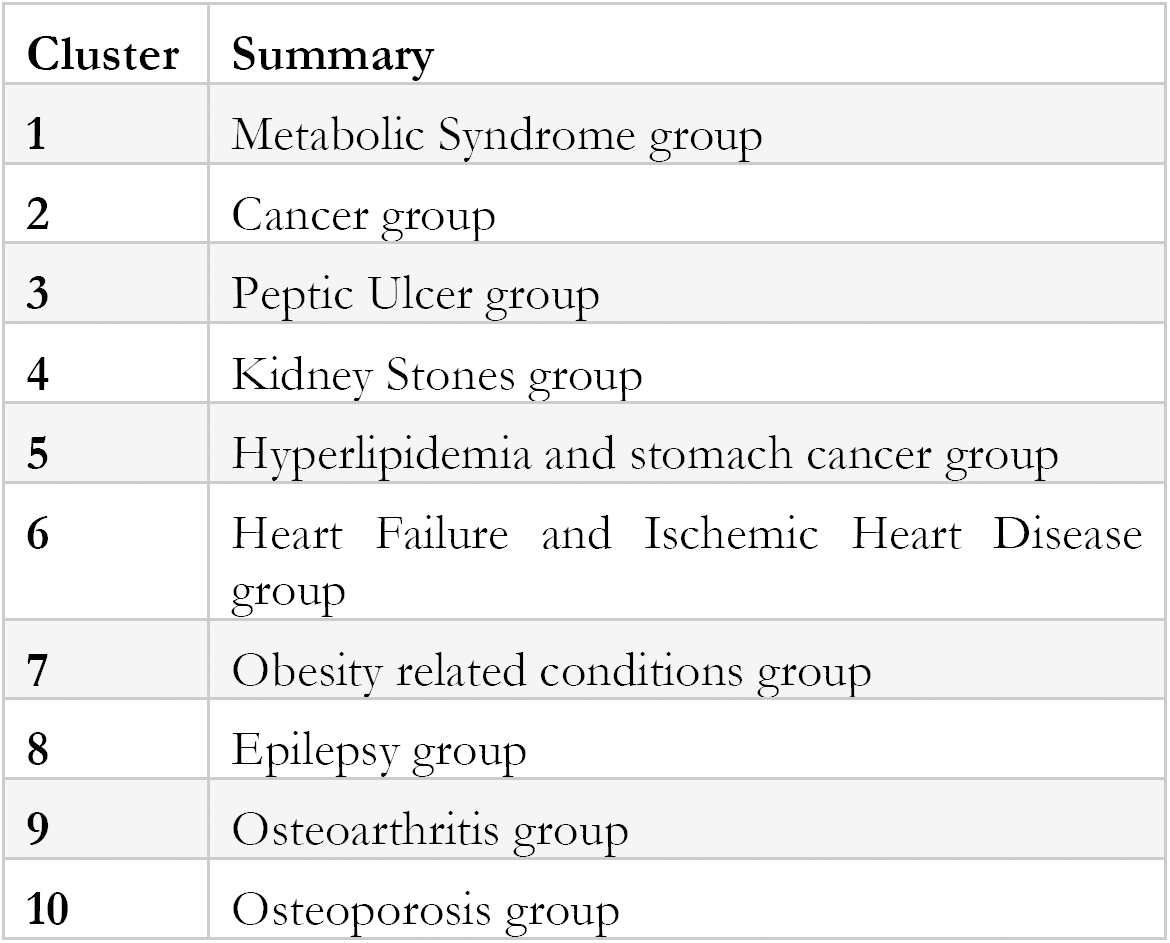
A summary of important clinical findings from each cluster.

## 5. DISCUSSION

In the present study we use the HAC approach to categorize cohorts of multimorbid patient with analogous multimorbidities in the Ohio population. A clinically relevant summarization of the comorbid conditions in every cohort is presented (Table 2).

Our first cluster had 1,193 members. It enclosed members with an average age 68 years, with more females than males (females: 58%). This cohort over indexed for oral cancer (87%) and metabolic syndrome (23%). Indeed, a 2015 meta-analysis revealed an association between metabolic syndrome and oral cancer^23^. The 2^nd^ cohort over indexed in females (55% females) with a lower age (median 62.2 years) with disorders like ovarian cancer (5.1%), cystic fibrosis (1.7%), pancreatic cancer (2.1%), hepatitis (6.9%) and post-partum birth disorders (0.2%). This was also our youngest cluster. Recent research has shown a linkage between ovarian cancers, pancreatic cancer and post-partum disorders further strengthening these causal findings^24–26^.

Cluster three consisted of males (58%) with a higher median age (age 75.7 years). This cohort over indexed on stomach conditions like peptic ulcers (82%), cholecystitis (99%), non-specific gastric dyspepsia (79%). Interestingly, we also found the highest incidence of lung cancer (23%) in this patient cohort. This association has previously been attributed to metastasis^27^.

Cluster 4 was gender equal and contained a high proportion of older patients with (median age 76.4 years; 51 % females; 49% male) with 92% of the members with kidney stones. Other conditions also seen in this cluster included colorectal cancer (4%), hypertension (67%), multiple sclerosis (13%) and congenital heart disease (2%). Alexander and colleagues, in a retrospective study have previously shown the how chronic hypertension is strongly linked and indicative of calcium oxalate crystal formation hinting at the root cause of this association^28^. Cluster 5, contained 1139 patients and a very high proportion of patients with hyperlipidemia (90%) and stomach cancer (4%). Recent research has shown significant association between the two conditions^29,30^.

Cluster 6 contained 1,494 patients, was more male skewed (57%), with a median age of 72.9 years. This cluster was predominantly dominated by heart diseases like heat failure (0.7%) and ischemic heart disease (0.4%).

Cluster 7 also over indexed in females (67% females) with a lower age (median 69.9 years) who reported a myriad of related disorders including chronic thyroid conditions (2.9%), endometrial cancer (3.3%), female infertility (0.2%), disrupted childhood disorder (0.2%) and obesity (24.8%). Comparable co/multimorbid relations have widely been considered in these patientis^31,32^.

Cluster 8, had a comparatively higher median age (age 77.5 years) with the highest index of males compared to females in the population (59.3% males) with a bulk of the members reporting epilepsy (78%). Epilepsy, one of the most prevalent neurological conditions with no known cure, is frequently seen in this subgroup^33^. Cluster 9 had more females (52%) with a median age of 79.2 years with a high proportion of patients with osteoarthritis (37%) and low vision blindness (52%). Cluster 10, our smallest cluster with only 507 members, was also our oldest with a median age of 81.1 years and a higher proportion of females (57%). This cluster consisted of diseases like osteoporosis (87%) and, rheumatoid arthritis (71%) and diverticular disease (64%). Osteoporosis and rheumatoid arthritis have often been linked to have similar mechanisms linked to interleukin-6 dysfunction ^34–36^ and this association further strengthens this hypothesis. This study has identified clinically relevant multimorbid clusters in a general population and can thus be used to profile patients in other geographies.

## BIBLIOGRAPHY

1. Triposkiadis, F. et al. Reframing the association and significance of co-morbidities in heart failure. Eur. J. Heart Fail. 18, 744–758 (2016).

2. Doos, L., Roberts, E. O., Corp, N. & Kadam, U. T. Multi-drug therapy in chronic condition multimorbidity: a systematic review. Fam. Pract. 31, 654–663 (2014).

3. Xu, X., Mishra, G. D. & Jones, M. Evidence on multimorbidity from definition to intervention: An overview of systematic reviews. Ageing Res. Rev. 37, 53–68 (2017).

4. Smith, S. M., Wallace, E., O’Dowd, T. & Fortin, M. Interventions for improving outcomes in patients with multimorbidity in primary care and community settings. Cochrane Database Syst. Rev. 3, CD006560 (2016).

5. Bratzke, L. C. et al. Self-management priority setting and decision-making in adults with multimorbidity: a narrative review of literature. Int. J. Nurs. Stud. 52, 744–755 (2015).

6. Singh, S. P., Karkare, S., Baswan, S. M. & Singh, V. P. Agglomerative Hierarchical Clustering Analysis of co/multi-morbidities. 18

7. Kavuri, V. C. & Liu, H. Hierarchical clustering method to improve transrectal ultrasound-guided diffuse optical tomography for prostate cancer imaging. Acad. Radiol. 21, 250–262 (2014).

8. Stryhn, H. & Christensen, J. The analysis-hierarchical models: past, present and future. Prev. Vet. Med. 113, 304–312 (2014).

9. Peña-Malavera, A., Bruno, C., Fernandez, E. & Balzarini, M. Comparison of algorithms to infer genetic population structure from unlinked molecular markers. Stat. Appl. Genet. Mol. Biol. 13, 391–402 (2014).

10. Adewale, A. J. et al. Understanding hierarchical linear models: applications in nursing research. Nurs. Res. 56, S40–46 (2007).

11. Cohen-Addad, V., Kanade, V., Mallmann-Trenn, F. & Mathieu, C. Hierarchical Clustering: Objective Functions and Algorithms. ArXiv170402147 Cs (2017).

12. Gollub, J. & Sherlock, G. Clustering microarray data. Methods Enzymol. 411, 194–213 (2006).

13. WHO | International Classification of Diseases, 11th Revision (ICD-11). WHO Available at: http://www.who.int/classifications/icd/en/. (Accessed: 19th June 2018)

14. Singh, S. P. Quantitative analysis on the origins of morphologically abnormal cells in temporal lobe epilepsy. (University of Cincinnati, 2015).

15. Singh, S. P. Advances in Epilepsy: A data science perspective. Data Science Journal 58, 89–92 (2016).

16. Singh, S. P. & Karkare, S. Stress, Depression and Neuroplasticity. arXi. eprint arXiv:1711.09536 (2017). doi:arXiv:1711.09536

17. Singh, S. P., Karkare, S., Baswan, S. M. & Singh, V. P. The Application of Text Mining Algorithms In Summarizing Trends in Anti-Epileptic Drug Research. Int. J. Stat. Probab. 7, 11 (2018).

18. Singh, S. P. & Karkare, S. 10K Pubmed Abstracts related to AntiEpileptic Drugs. (2018). doi:10.6084/m9.figshare.5764524.v1

19. Singh, S. P. & Singh, V. P. Quantitative Analysis on the role of Raffinose Synthase in Hippocampal Neurons. bioRxiv (2017). doi:10.1101/240192

20. Singh, S. P., Singh, S. P., Fatima, N., Kubo, E. & Singh, D. P. Peroxiredoxin 6-A novel antioxidant neuroprotective agent. Neurology 70, A480–A481 (2008).

21. Gower, J. C. A General Coefficient of Similarity and Some of Its Properties. Biometrics 27, 857 (1971).

22. Dawson, K. J. & Belkhir, K. An agglomerative hierarchical approach to visualization in Bayesian clustering problems. Heredity 103, 32–45 (2009).

23. Chang, C.-C. et al. Metabolic syndrome and health-related behaviours associated with pre-oral cancerous lesions among adults aged 20-80 years in Yunlin County, Taiwan: a cross-sectional study. BMJ Open 5, e008788 (2015).

24. Pittman, M. E., Brosens, L. A. A. & Wood, L. D. Genetic Syndromes with Pancreatic Manifestations. Surg. Pathol. Clin. 9, 705–715 (2016).

25. Xu, J. et al. Adenovirus-mediated overexpression of cystic fibrosis transmembrane conductance regulator enhances invasiveness and motility of serous ovarian cancer cells. Mol. Med. Rep. 13, 265–272 (2016).

26. Zagouri, F., Dimitrakakis, C., Marinopoulos, S., Tsigginou, A. & Dimopoulos, M.-A. Cancer in pregnancy: disentangling treatment modalities. ESMO Open 1, e000016 (2016).

27. Miyazaki, J., Hirota, S. & Abe, T. Metastasis of lung cancer to the gastrointestinal tract, presenting with a volcano-like ulcerated mass. Dig. Endosc. Off. J. Jpn. Gastroenterol. Endosc. Soc. 27, 397–398 (2015).

28. Alexander, R. T. et al. Thiazide Diuretic Dose and Risk of Kidney Stones in Older Adults: A Retrospective Cohort Study. Can. J. Kidney Health Dis. 5, 2054358118787480 (2018).

29. Byard, R. W. The complex spectrum of forensic issues arising from obesity. Forensic Sci. Med. Pathol. 8, 402–413 (2012).

30. Singh, P. P. & Singh, S. Statins are associated with reduced risk of gastric cancer: a systematic review and meta-analysis. Ann. Oncol. Off. J. Eur. Soc. Med. Oncol. 24, 1721–1730 (2013).

31. Hanley, P., Lord, K. & Bauer, A. J. Thyroid Disorders in Children and Adolescents: A Review. JAMA Pediatr. 170, 1008–1019 (2016).

32. Bagur, J. et al. Psychiatric disorders in 130 survivors of childhood cancer: preliminary results of a semi-standardized interview. Pediatr. Blood Cancer 62, 847–853 (2015).

33. American Epilepsy Society | Working Toward a World without Epilepsy. Available at: https://www.aesnet.org/. (Accessed: 25th November 2017)

34. Hoes, J. N., Bultink, I. E. M. & Lems, W. F. Management of osteoporosis in rheumatoid arthritis patients. Expert Opin. Pharmacother. 16, 559–571 (2015).

35. Clayton, E. S. & Hochberg, M. C. Osteoporosis and osteoarthritis, rheumatoid arthritis and spondylarthropathies. Curr. Osteoporos. Rep. 11, 257–262 (2013).

36. Matuszewska, A. & Szechiński, J. [Mechanisms of osteoporosis development in patients with rheumatoid arthritis]. Postepy Hig. Med. Doswiadczalnej Online 68, 145–152 (2014).

